# ZmSWEET Sucrose transporters expressed in the endosperm adjacent to the maize embryo are necessary for carbon partitioning and embryo growth

**DOI:** 10.64898/2026.02.24.707659

**Authors:** Yannick Fierlej, Laurine Grazer, Abdelsabour G.A. Khaled, Matthias Langer, Emilie Montes, Thibaut Perez, Laura Gallo, Benoît Lacombe, Philippe Nacry, Hélène Duplus-Bottin, Nicolas M. Doll, Ljudmilla Borisjuk, Hardy Rolletschek, Gwyneth Ingram, Peter M. Rogowsky, Thomas Widiez

**Affiliations:** Laboratoire Reproduction et Développement des Plantes, ENS de Lyon, Lyon 1 Université, CNRS, INRAE, F-69342 Lyon, France; Department of Genetics, Faculty of Agriculture, Sohag University, 82786 Sohag, Egypt; Leibniz-Institute of Plant Genetics and Crop Plant Research (IPK), Corrensstrasse 3, 06466 Seeland Gatersleben, Germany; IPSiM, Université de Montpellier, CNRS, INRAE, Institut Agro, Montpellier Cedex, France; Laboratory of Biology and Modeling of the Cell, Universite de Lyon, Ecole Normale Superieure de Lyon, CNRS, Universite Claude Bernard Lyon 1, Lyon, France

**Keywords:** Sugar, transport, SWEET, EAS, embryo, endosperm, Zea mays

## Abstract

In cereals such as maize, the kernel accumulates large quantities of storage compounds, including carbohydrates, lipids, and proteins, a process that requires tight regulation of nutrient transport. Seeds are composed of distinct tissues: the embryo, the endosperm, and maternal tissues that are symplastically isolated (not connected through plasmodesmata), necessitating specialized nutrient transfer mechanisms. In maize, nutrient transfer from maternal tissues to the endosperm via specialized basal endosperm transfer layer (BETL) cells is well characterized. However, nutrient transfer at the endosperm/embryo interface remains poorly understood. Consequently, the routes by which maternal carbon-derived sugars support embryo growth are still unclear. Our previous transcriptomic profiling uncovered a novel Endosperm domain Adjacent to the embryo Scutellum (EAS) with strong enrichment for transporter genes. Notably, genes encoding three sugar transporters from the SWEET (Sugars Will Eventually be Exported Transporters) family are highly and preferentially expressed in the EAS, suggesting the existence of a specialized sugar transfer mechanism at this interface. We show that the ZmSWEET proteins encoded by these genes are membrane-localized sucrose transporters and are functionally important for kernel development. A gene-edited triple *zmsweet14a/14b/15a* knock-out mutant exhibits reduced kernel weight and embryo size, significantly decreased embryo oil accumulation at maturity, and altered carbon partitioning within the kernel. In addition to these defects, mutant kernels display a significant reduction in primary root length during germination, indicating either lasting physiological consequences of disrupted sucrose transport during seed development or an additional role for these SWEET transporters during germination. Together, our findings demonstrate that sucrose transport at the endosperm/embryo interface is critical for proper carbon allocation, embryo development, and seed vigor, and identify the EAS as a key functional domain and potential target for improving seed composition.

## Introduction

Seeds play a central role in the global food system, serving as a major source of calories through both direct human consumption and indirect use in animal feed. This essential contribution to global energy intake stems from the fact that seeds store reserves, primarily carbohydrates, proteins, and lipids. Seed are complex biological system formed of three main tissues: the embryo, the endosperm and maternal tissues. The embryo is fully surrounded by the endosperm, and both of them are further enclosed within the maternal tissues (Lafon-Placette and Köhler, 2014; Widiez *et al*., 2017; Doll and Ingram, 2022). Plant species exhibit substantial variation in both seed size and the composition of these storage products (Sreenivasulu and Wobus, 2013). In maize for example, the endosperm remains persistent and serves as a starch reservoir, while the embryo accumulates high concentrations of oil (Dochlert, 1990). Thus, seeds represent a heterotrophic “sink” tissue and rely on an important organic carbon supply from the photoautotrophic source tissues, especially in maize (Palmer *et al*., 1973). Sucrose is the primary photosynthate transported predominantly from mature photosynthetic leaves via the vascular system to non-photosynthetic sink tissues, where it supports their growth and development (Gifford and Evans, 1981; Koch, 2004; Ruan, 2014).

Upon arrival at the proximity of developing seeds, sucrose exits the vascular system at the phloem terminus through a process known as phloem unloading (Thorne, 1985; Patrick and Offler, 2001; Braun, 2022). Due to the organization of the seed, the maternal tissues are symplastically isolated from the endosperm cells, meaning that their cytoplasms are not connected via plasmodesmatal pore (Stadler *et al*., 2005; Widiez *et al*., 2017). Consequently, sugar coming from the maternal vascular tissues must go through the apoplast and be loaded within the endosperm by crossing plasma membranes (Patrick and Offler, 2001; Povilus and Gehring, 2022). In maize, the basal endosperm transfer layer (BETL), contains cell types specialized in nutrient transport from the mother plant to the endosperm, designed for nutrient uptake, and can be viewed together with the placento-chalazal cells as an important interface regulating the sugar entrance from the sporophyte to the seed (Chourey and Hueros, 2017; Povilus and Gehring, 2022). This maternal-to-filial nutrition transfer is central to grain development and yield, as illustrated by the severe shrinkage of maize kernels when nutrition-related genes are mutated (Kang *et al*., 2009; Sosso *et al*., 2015; Yang *et al*., 2022). Similar apoplastic sugar loading should occur at the endosperm-embryo interface (Doll and Ingram, 2022). The nutritional importance of maize endosperm-embryo interfaces is indirectly highlighted by the expression of sugar transporters as well as an invertase inhibitor in both the embryo-surrounding region of the endosperm (ESR, 4-10 DAP) and later in development (9-20 DAP) in the endosperm adjacent to scutellum (EAS) interface (Bate *et al*., 2004; Doll *et al*., 2020; Shen *et al*., 2022). So far, our understanding of the mechanisms underlying apoplasmic pathways for carbohydrates transport across maize filial interfaces (embryo/endosperm) remains partial.

Efficient carbohydrate partitioning requires the specific activity of membrane sugar transporters. Sugar transporters are essential components of both intracellular trafficking and apoplastic transport pathways and form three key families in plants: MSTs (monosaccharide transporters), SUTs (sucrose transporters) and SWEETs (Sugars Will Eventually be Exported Transporters) (Eom *et al*., 2015; Keller and Neuhaus, 2025). MSTs and SUTs belong to the major facilitator superfamily (MFS), have twelve transmembrane domains, and most characterized members function as active transport systems using the driving force of the H+ ATPase pump (Lalonde *et al*., 2004; Niño-González *et al*., 2019; Geiger, 2020). SWEETs are a class of uniporters characterized by 7 transmembrane domains that transport hexose and sucrose and mediate sugar influx or efflux along a concentration gradient (Chen *et al*., 2015a; Anjali *et al*., 2020). In *Arabidopsis thaliana*, a cascade of sequentially expressed SWEET sucrose transporters in the seed coat and endosperm provides nutrition for the embryo. The *atsweet11;12;15* triple mutant has a wrinkled seed phenotype, with accumulation of starch in the seed coat and delayed embryonic development (Chen *et al*., 2015b). Crucial role of SWEETs in seed physiology has been demonstrated in several crops. In soybean (*Glycine max*), GmSWEET15a and GmSWEET15b are essential for early stages of seed development by mediating sucrose export from the endosperm towards the embryo. Knock-out of both genes causes retarded embryo growth, endosperm persistence and seed abortion (Wang *et al*., 2019). In tomato (*Solanum lycopersicum*), knock-out of *SlSWEET15* caused defects in seed filling and embryo development, suggesting that SlSWEET15 is required for sucrose unloading from the seed coat to the growing embryo (Ko *et al*., 2021). In rice (*Oryza sativa*), OsSWEET11, OsSWEET14 and OsSWEET15 have been shown to mediate sucrose efflux in developing caryopses and play key roles in grain filling. *ossweet11;14* and *ossweet11;15* double mutants both exhibited reduced grain weight and abnormal starch accumulation in the pericarp (Yang *et al*., 2018; Fei *et al*., 2021; Hu *et al*., 2023). In maize, the BETL expressed ZmSWEET4c is responsible for transferring hexoses (glucose and fructose) of developing kernels from the maternal tissues to the endosperm. Knock-out mutation of *ZmSWEET4c* generates a dramatic reduction of endosperm size and kernel weight (Sosso *et al*., 2015). In addition to ZmSWEET transporter, mutation of maize sugar NPF transporter gene Sucrose and Glucose Carrier 1 (ZmSUGCAR1/ZmNPF7.9), which is expressed in BETL interface, also cause impairment of endosperm enlargement resulting in poorly filled kernels with empty pericarp (Yang *et al*., 2022). These data on maize illustrate the key importance of BETL region for sugar entry into the maize seed (Doll *et al*., 2017).

Our previous work identified a novel endosperm cell type adjacent to the embryo, termed the Endosperm Adjacent to Scutellum (EAS) (Doll *et al*., 2020). EAS cell layers represent the largest endosperm surface in direct contact with the embryo and are progressively eliminated through cell death, thereby creating space for embryo growth (Doll *et al*., 2025). Notably, the EAS cells exhibit enriched expression of three SWEET transporters (ZmSWEET14a, ZmSWEET14b, and ZmSWEET15a) (Doll *et al*., 2020), providing an opportunity to investigate the role of sugar transport at this endosperm–embryo interface. In the present study, we demonstrate that these SWEETs play a critical role in kernel development, embryo growth and carbon partitioning within the maize kernel.

## Results

### Three plasma-membrane localized clade III SWEET sucrose transporters show preferential expression in the EAS endosperm sub-domain

Protein sequences of the SWEET family transporters from maize, rice and Arabidopsis were used to construct a phylogenetic tree (**Figure 1A**). The three ZmSWEET encoding genes (*ZmSWEET14a* = Zm00001eb113080; ZmSWEET14b = Zm00001eb170150; ZmSWEET15a = Zm00001eb180830), previously identified as showing enriched expression in the EAS at 13 DAP (Doll *et al*., 2020) were found to belong to clade III of SWEET family (**Figure 1A**), as previously reported (Wang *et al*., 2024). Comparative expression profiles revealed that *ZmSWEET15a* is more strongly expressed in the EAS compared to *ZmSWEET14a* and *ZmSWEET14b* (Doll *et al*., 2020) (**Supplemental figure S1A**). Expression profiles across different maize tissues also show expression of *ZmSWEET14a* and *ZmSWEET14b* in primary roots during germination, and expression of *ZmSWEET15a* in meiotic tassels and leaf bases (Hoopes *et al*., 2019; Zhu *et al*., 2022) (**Supplemental figure S1B**). Since clade III of SWEET family are known to be involved in sucrose transport (Eom *et al*., 2015), a yeast heterologous system was used to assess whether ZmSWEET14a, ZmSWEET14b and ZmSWEET15a are able to transport sucrose. Expression of ZmSWEET14b or ZmSWEET15a allows the yeast mutant strain SUSY7, which enables phenotypic recognition of a sucrose carrier activity (Riesmeier *et al*., 1992), to grow more efficiently using sucrose as the sole carbon source, indicating that ZmSWEET14b and ZmSWEET15a can transport sucrose in this system (**Figure 1B**). Consistent with a role in intercellular sucrose transport, subcellular localization analyses in maize mesophyll protoplasts indicated that ZmSWEET15a:mCitrine co-localizes with the plasma membrane marker protein LTI6b:mTurquoise (**Figure 1C**).

**Figure 1.**
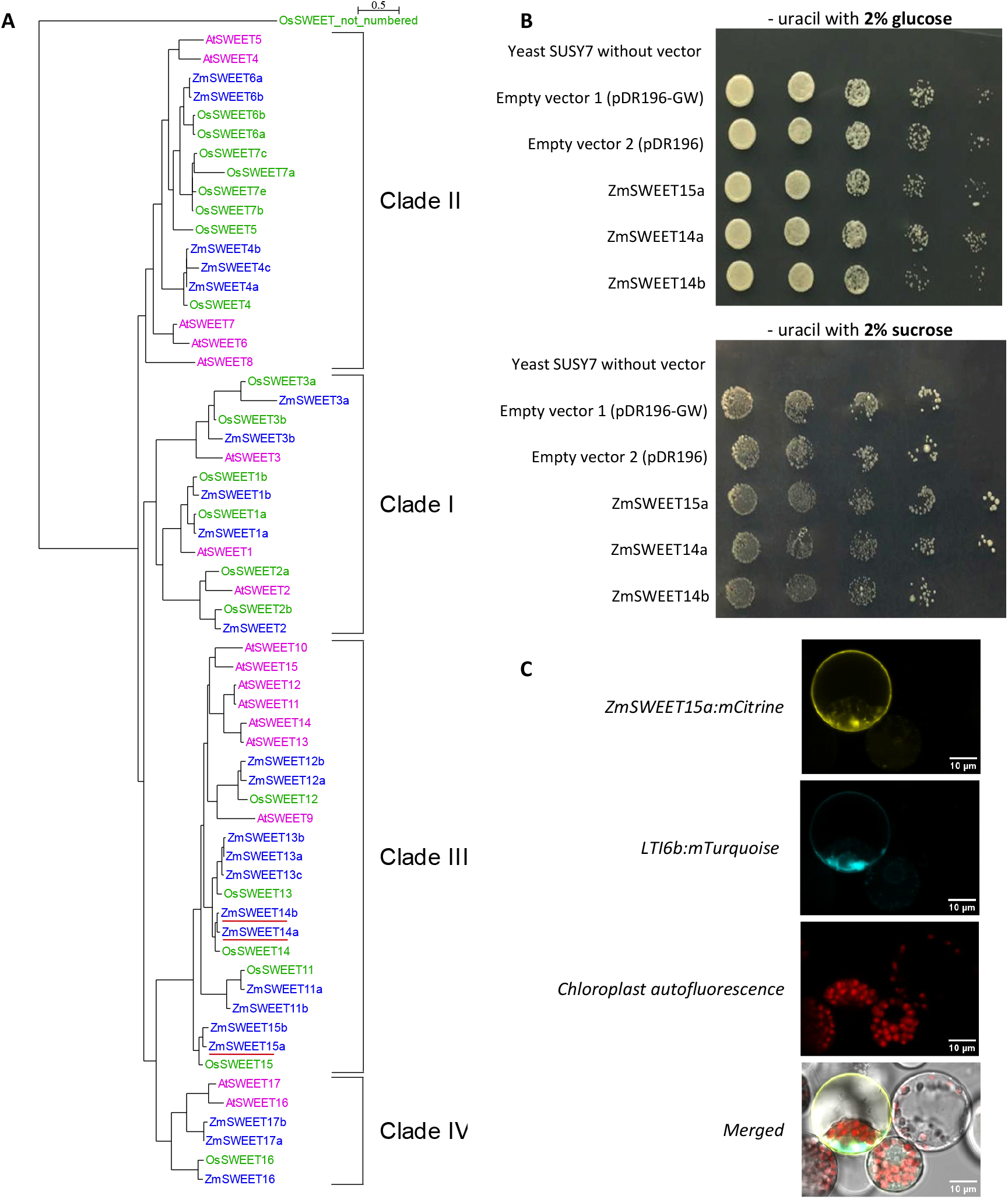
Three ZmSWEET 14a/14b/15a are sucrose transporter which are expressed in the EAS region and localize to the plasma membrane in protoplast. (**A**) Phylogenetic analysis of SWEET transporters from *Zea mays* (Zm, blue), *Oryza sativa* (Os, green), and *Arabidopsis thaliana* (At, magenta). Three *ZmSWEET* genes from Clade III, showing preferential expression in the endosperm adjacent to the scutellum (EAS), are highlighted with a red underline. (**B**) Heterologous expression of *ZmSWEET14a*; *ZmSWEET14b* and *ZmSWEET15a* in the SUSY7 yeast strain which allows the phenotypic recognition of a sucrose carrier activity. Glucose containing medium is used as control. (**C**) Subcellular localization of ZmSWEET15a:mCitrine in maize protoplasts. The *LTI6b:mTurquoise* was used as plasma membrane marker. Fluorescent signals from mCitrine, mTurquoise, and chloroplast autofluorescence were visualized by confocal laser scanning microscopy.

### Embryos of the *zmsweet14a/14b/15a* triple mutant are smaller and accumulate less oil at maturity

To investigate the function of ZmSWEET14a, ZmSWEET14b and ZmSWEET15a in kernel development, a CRISPR/Cas9-mediated genome editing approach was used to induce mutations. Two single guide RNAs (sgRNAs) were designed to target fully identical sequences in exon 4 of *ZmSWEET14a* and *ZmSWEET14b*, while two additional sgRNAs were designed to target *ZmSWEET15a* (**Fig. 2A-B**). Following maize transformation, gene-specific primer pairs (**Fig. 2A**) were used to detect sequence alterations at the CRISPR/Cas9 target sites through Sanger sequencing. This analysis revealed short insertions and deletions (indels), resulting in three distinct mutant alleles for both *ZmSWEET14a* and *ZmSWEET14b*, and one mutant allele for *ZmSWEET15a* (**Fig. 2B**). All identified alleles, except *zmsweet14b_2*, are frameshift mutations that disrupt the second conserved MtN3_slv (<u>PFAM accession number: PF03083</u>) family protein domain and create premature stop codons (**Fig. 2C**). Crosses were performed to remove the CRISPR/Cas9 T-DNA cassette and generate a *zmsweet14a_1/zmsweet14b_3/ zmsweet15a_1* triple mutant subsequently referred to as *zmsweet14a/14b/15a* for simplicity. To investigate morphological characteristics of the embryo and to visualize oil distribution in mature kernels, nuclear magnetic resonance imaging (MRI) was employed as non-destructive and non-invasive technique to acquire three-dimensional images of intact embryos from *zmsweet14a/14b/15a* mutants and WT (**Fig. 3**). This analysis revealed that embryos of the *zmsweet14a/14b/15a* mutant were significantly (∼16 %) smaller in volume compared to WT, although their overall shape remained similar (**Fig. 3 A-D, I**). Measuring the kernel weight also revealed a reduced seed weight for *zmsweet14a/14b/15a* mutant as compared to wild-type (**Supplemental figure S2-A**). Storage oils (triacylglycerides) constitute the primary dry mass component of the embryo and can be visualized via MRI. Oil distribution showed spatial heterogeneity in both WT and *zmsweet14a/14b/15a* embryos (**Fig. 3 E-H**). The highest lipid concentrations were consistently observed in the scutellum, with lower levels in the radicle and plumule regions. This pattern is typical for maize embryos and was found in both genotypes. However, the total oil content per embryo was significantly reduced (by ∼21%) in the mutant compared to WT (**fig. 3J**). When comparing the average lipid signal per unit volume (the relative concentration of oil within the seed volume), no statistically significant difference was found between genotypes (**Supplemental figure S2B, C**), which can be explained by a proportional reduction in both embryo size and total oil accumulation in the mutant. Thus, this indicates that the reduced oil content in the *zmsweet14a/14b/15a* mutant embryo is mainly due to its smaller size rather than a metabolic defect in oil accumulation in the embryo.

**Figure 2.**
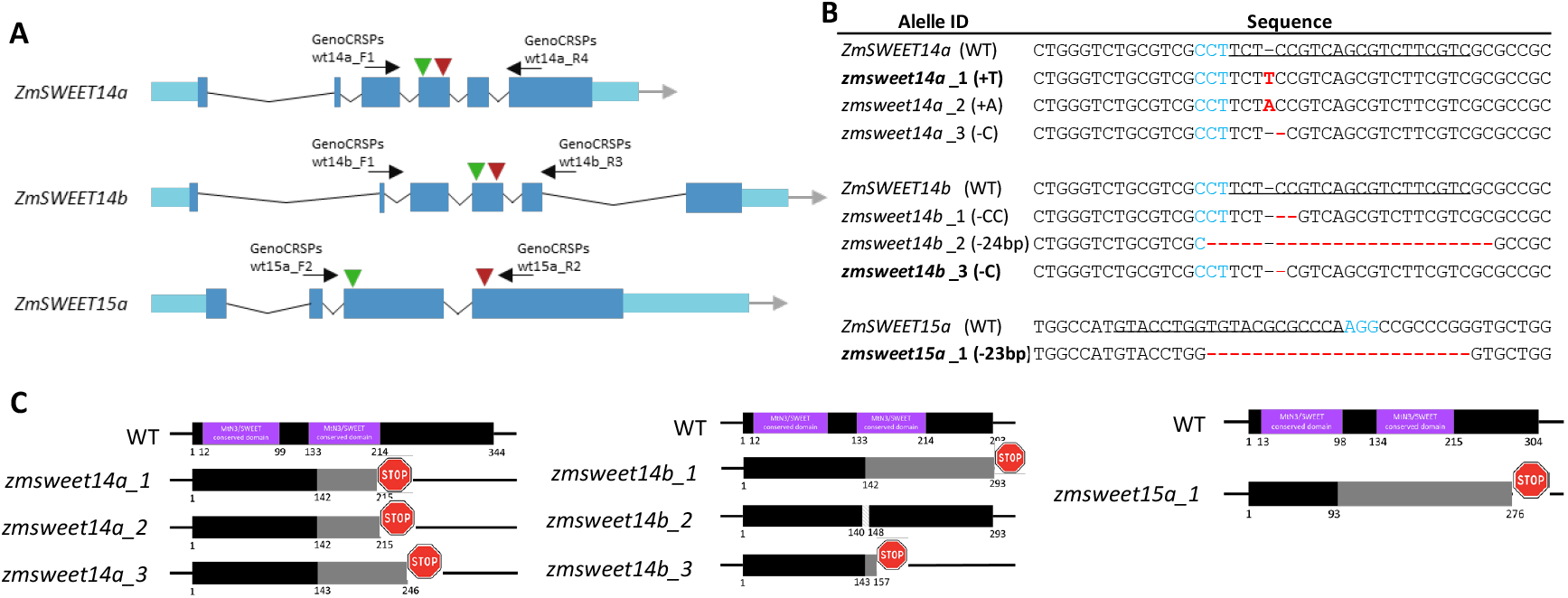
Generation of knock out mutants in *ZmSWEET14a, ZmSWEET14b* and *ZmSWEET15a* by genome editing. (**A**) Schematic representation of the gene models for *ZmSWEET14a, ZmSWEET14b*, and *ZmSWEET15a*. Light blue boxes indicate untranslated regions (UTRs), blue boxes represent exons, and thin black lines indicate introns. Green and red triangles mark the positions of sgRNA1 and sgRNA2, respectively, used for CRISPR-Cas9 genome editing. Black arrows indicate the locations of primers used for genotyping and mutation characterization. (**B**) Overview of the mutant alleles obtained. DNA sequences targeted by sgRNA are underlined. PAM sequences are shown in blue. Letters in red and red hyphens indicate insertions and deletions respectively. Alleles highlighted in bold were selected to generate the triple mutant line. (**C**) Predicted consequences of the mutations at the protein level. Purple boxes indicate conserved MtN3/SWEET domains. Black boxes represent in-frame amino acids, while grey boxes indicate out-of-frame sequences. Red STOP symbols denote the positions of premature stop codons.

**Figure 3.**
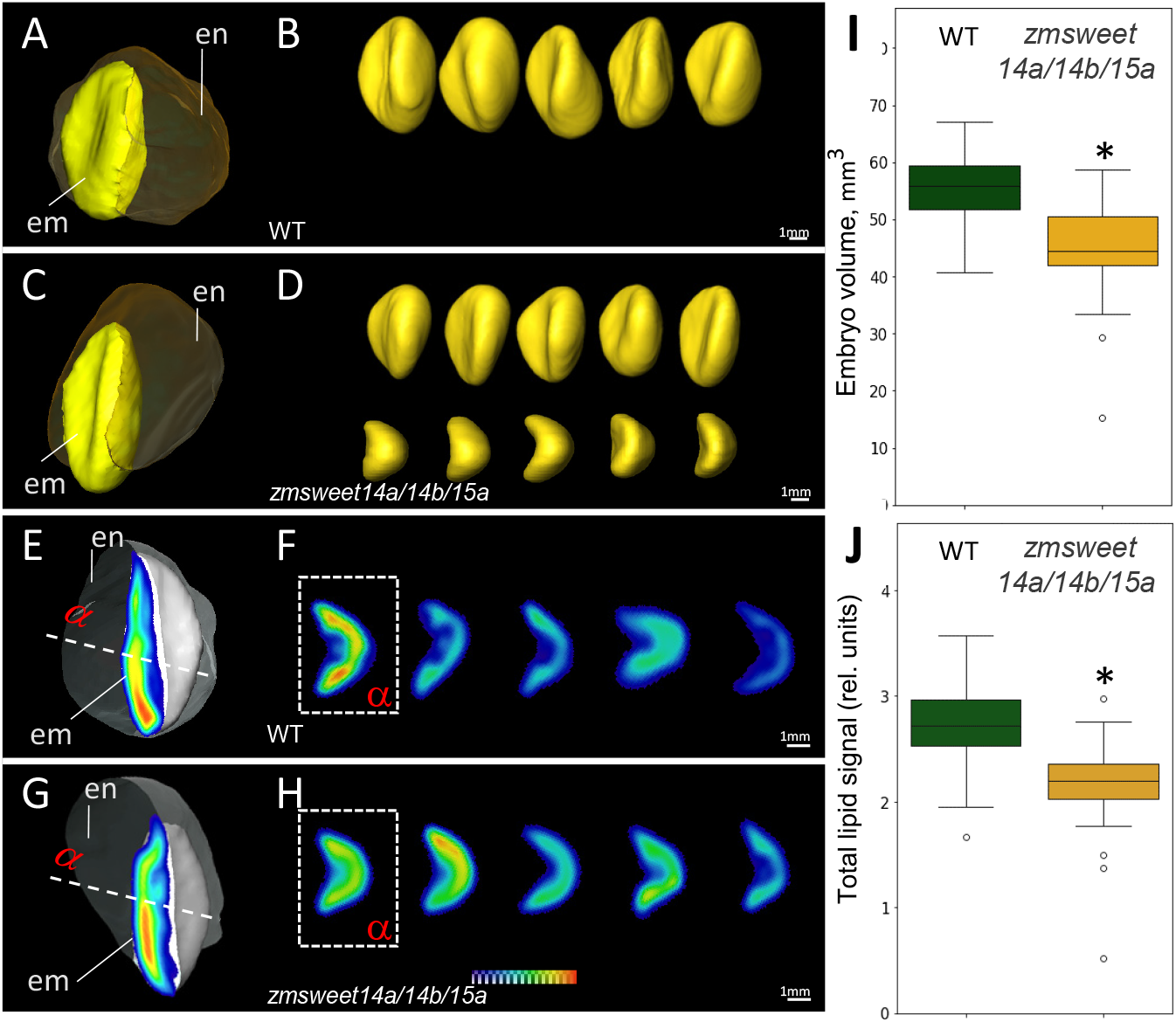
Comparative magnetic resonance imaging of WT and mutant *zmsweet14a/14b/15a* kernels at maturity. (**A,B**) Representative MRI image showing three-dimensional wild-type (WT) maize kernels (**A**) and individual intact embryos (sagital and axial view) (**B**); Images shown are representative of five biological replicates. All images were digitally extracted for comparison and are displayed at the same scale. (**C,D**) Equivalent images for *zmsweet14a/14b/15a* mutant kernels (**C**) and embryos (**D**). (**E,F**) Distribution of lipid-specific signal in embryonic tissues of WT embryos (**E**): virtual sections (axial) through the embryos (**F**); signal intensity is colour-coded in relative units. (**G,H**) Same for *zmsweet14a/14b/15a* mutant embryos. (**I**) Quantification of embryo volume in WT and *zmsweet14a/14b/15a* mutant embryos. (**J**) Quantification of total lipid signal per embryo in WT vs. *zmsweet14a/14b/15a* mutant. Data are given in relative units. Asterisks in panels **(I)** and **(J)** indicate statistically significant differences (p<0.05, n=40, T-test). Abbreviations: en, endosperm; em, embryo.

### The triple *zmsweet14a/14b/15a* mutant is impaired in germination and seedling establishment

To assess the germination capacity and seedling development of the *zmsweet14a/14b/15a* triple mutant, two different assays were conducted. First, the rolled towel method (Gonzales *et al*., 2024) was used by placing kernels in rolled-up germination paper. Primary root length was measured at 6 and 9 days after sowing (DAS) (**Fig. 4A-B**). The *zmsweet14a/14b/15a* mutant exhibited a significant reduction in primary root length at both time points compared to WT (**Fig. 4B**). Secondly, a hydroponic assay was performed to evaluate multiple aspects of the root system architecture, as well as shoot and root fresh weight (**Fig. 4C-G**). At 11 DAS, *zmsweet14a/14b/15a* mutants showed a significant reduction in both shoot (∼27%) and root fresh weight (14%) as compared to WT (**Fig. 4D**). Additionally, the length of primary roots (∼16% inhibition) and the number of seminal roots (∼19% inhibition), but not their average length, were significantly reduced in the *zmsweet14a/14b/15a* mutant compared to WT (**Fig. 4E-F**). Taken together, these data suggest that the *zmsweet14a/14b/15a* mutation significantly impairs seedling root development.

**Figure 4.**
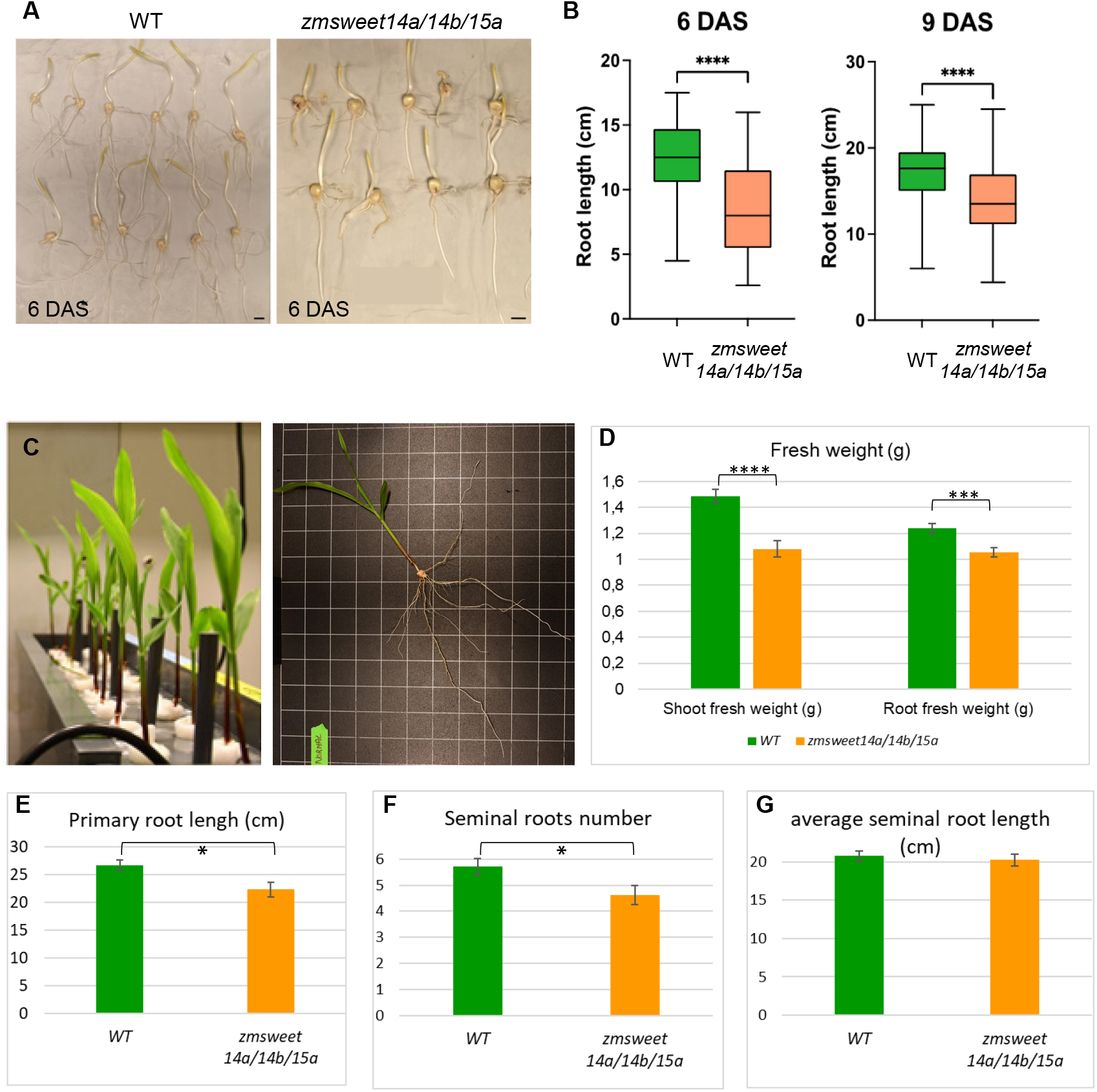
Triple mutant *zmsweet14a/14b/15a* are impaired in germinations Illustration of the germination assay on paper. (**B**) Quantification of primary root length at 6 and 9 days after sowing (DAS). Values represent the mean of 80–117 seedlings; error bars indicate standard error. (**C**) Illustration of experiment set-up for hydroponic assay. (**D**) Shoot and root fresh weigh, (**E**) primary root length, (**F**) seminal roots number and (**G**) average seminal root length of 11 DAS WT and triple *zmsweet14a/14b/15a* grown under hydroponic conditions. Values represent the mean of 13-20 seedlings; error bars indicate standard error. Asterisks denote statistical significance based on t-test: **** = *p* < 0.0001; *** = *p* < 0.001; * = *p* < 0.05.*15a*

### Carbon partitioning is altered in *zmsweet14a/14b/15a* maize kernels

Because *ZmSWEET14a, ZmSWEET14b* and *ZmSWEET15a* encode sucrose transporters with enriched expression in maize kernels, the influence of their combined loss of function was used to investigate their role in carbon partitioning during kernel development. To this end, the large size of the maize kernels was leveraged to examine the spatial distribution of sucrose by applying the Fourier-transform infrared (FTIR) protocol (Guendel *et al*., 2018). Sucrose distribution was successfully visualized in cryo-sectioned kernels at the early grain-filling stage (13 days after pollination, DAP) and later at 17 DAP (**Fig. 5**). Analysis of wild-type (WT) kernel sections revealed an apical–basal sucrose gradient, with higher sucrose accumulation in the basal endosperm compared to the upper endosperm (**Fig. 5C, E**). Strong sucrose accumulation was also observed in maternal tissues, notably in the pedicel region where vascular strands terminate and assimilate unloading occurs, at the apical tip of the pericarp, and in a narrow strip of pericarp adjacent to the embryo (**Fig. 5C, E**). Comparison between WT and *zmsweet14a/14b/15a* mutant kernels showed that the overall sucrose distribution pattern remained largely unchanged. However, a trend toward reduced sucrose accumulation was detected in the basal endosperm at both 13 and 17 DAP, and in the embryo at 13 DAP in the mutant background (**Fig. 5D, F**). Given the semi-quantitative nature of FTIR imaging, a stable isotope tracing assay was developed to complement these observations. This involved feeding 13-DAP kernels with uniformly labeled ^13^C-sucrose, in which all carbon atoms in the sucrose backbone are ^13^carbon (^13^C) (**Fig. 6A**). The ^13^C accumulation of different tissues was measured using isotope-ratio mass spectrometry (IRMS) in dissected kernel compartments: the embryo, and the basal and upper parts of the endosperm. The two endosperm regions were defined by a horizontal line drawn at the top of the embryo: endosperm tissue below this line was considered as basal endosperm, and tissue above as upper endosperm (**Fig. 6A**). **Figure 6B** shows ^13^C quantification in the different compartments after 24h, 48h and 72h of feeding with the ^13^C-sucrose solution. Regardless of the genotype, longer incubation times with the ^13^C-sucrose solution led to greater ^13^C accumulation in all compartments analyzed (**Fig. 6B**). Among the compartments, the basal endosperm accumulated the highest levels of ^13^C, followed by the upper endosperm and the embryo (**Fig. 6B**). This observation is consistent with the FTIR imaging, although the IRMS method measures total ^13^C, which could be present as ^13^C-sucrose but could also have been metabolized into various compound such as starch, amino acids or lipids for example. Focusing on the comparison between the WT and the *zmsweet14a/14b/15a* mutant, the triple mutant accumulated less ^13^C, with differences that became significant after 72h of incubation with the ^13^C-sucrose solution. Impairment of ^13^C accumulation was found in all three compartments of the *zmsweet14a/14b/15a* mutant, indicating a global disruption in carbon allocation due to the loss of these sucrose transporters in both the endosperm and the embryo.

**Figure 5.**
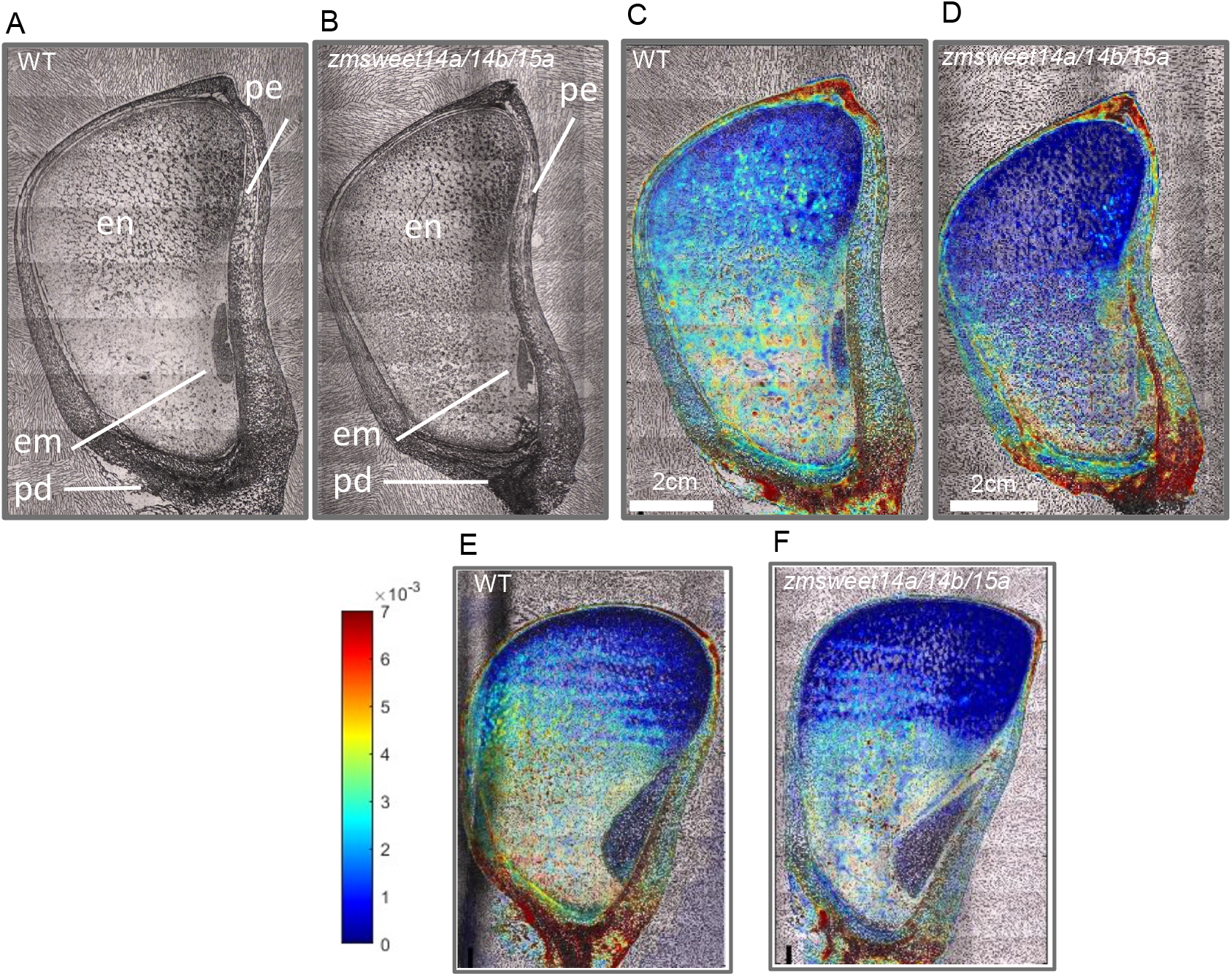
Imaging of sucrose distribution in the maize kernel of WT and *zmsweet14a/14b/15a* using FTIR microspectroscopy. **(A, B)** Bright-field images of cryo-sectioned kernels at 13 days after pollination (DAP) from wild-type (WT) (A) and *zmsweet14a/14b/15a* triple mutant (B). (C, D) Color map showing sucrose distribution across the same sections of WT (C) and mutant (D) kernels, as detected by FTIR microspectroscopy. (E,F) FTIR microspectroscopy on 17 DAP wild-type (WT) (E) and *zmsweet14a/14b/15a* triple mutant (F) kernel. Abbreviations: en – endosperm; em – embryo; pe – pericarp. Bar: 2 cm.

**Figure 6.**
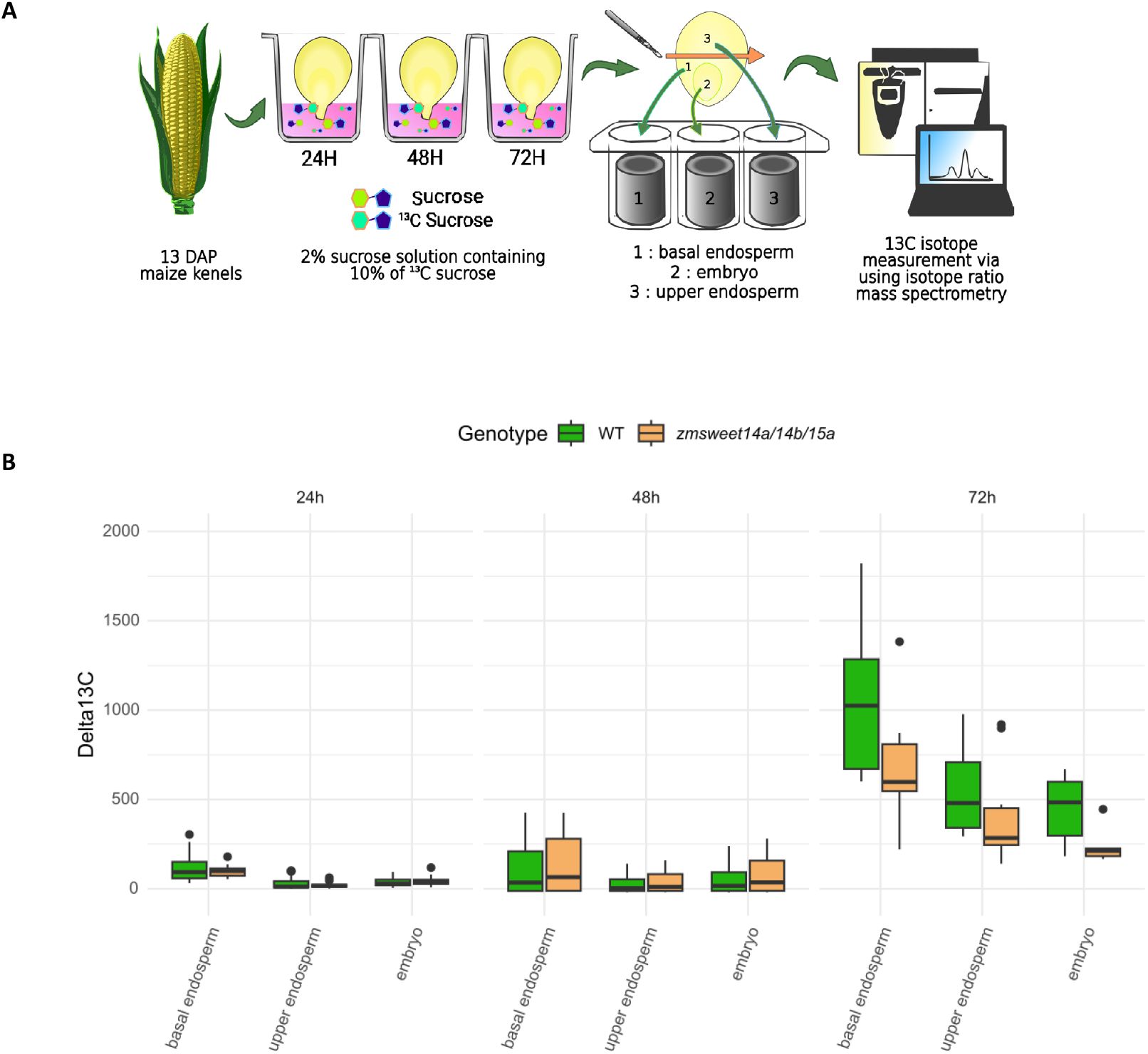
^13^C-sucrose feeding assay reveals altered carbon partitioning in the *zmsweet14a/14b/15a*. (**A**) Schematic representation of the ^13^C-sucrose feeding assay. Kernels at 13 days after pollination (DAP) from wild-type (WT) and *zmsweet14a/14b/15a* mutant plants were detached from the ear and immersed in a sucrose solution containing ^13^C-labeled sucrose for 24, 48, or 72 hours. After incubation, embryos and endosperms were dissected. The endosperm was further separated into basal and upper parts. (**B**) The ^13^C content derived from the labeled sucrose was quantified in each tissue using isotope ratio mass spectrometry in WT (green) and mutant (orange) kernels. Values represent the mean of 13-20 kernels.

## Discussion

### Apoplastic sugar transport and developmental outcomes at the embryo–endosperm interface

During seed development, nutrient fluxes must traverse two symplastic barriers: between maternal tissues and the endosperm, and between the endosperm and the embryo (Patrick and Offler, 2001). In maize apoplastic post-phloem transport to the endosperm BETL cells is facilitated by a suite of transporters and enzymes (Chourey and Hueros, 2017). Both sucrose and hexose transporters (Sosso *et al*., 2015; Yang *et al*., 2022) and cell wall invertases converting sucrose to hexoses (Kang *et al*., 2009) are critical for sugar transfer from maternal tissues to the endosperm at this interface. By contrast, much less is known about the mechanisms governing sugar transport at the endosperm–embryo interface. Our work focuses on the less-characterized maize endosperm-embryo interface, across which apoplastic sugar transport also occurs; the Endosperm Adjacent to the embryo Scutellum (EAS). Previous transcriptomic analyses identified *ZmSWEET14a, ZmSWEET14b* and *ZmSWEET15a* as being specifically expressed at this interface (Doll *et al*., 2020; Shen *et al*., 2022) providing a strong rationale to target them by genome editing. We show that mutation of these three EAS-expressed ZmSWEET sucrose transporters affects maize kernel and embryo size. A comparable expression pattern in the endosperm surrounding the embryo has been reported for the orthologous rice *OsSWEET15* and soybean *GmSWEET15* genes, with mutations leading to retarded embryo development, suggesting a conserved function for sugar transfer from the endosperm to embryo in diverse species (Wang *et al*., 2019; Hu *et al*., 2023). Soybean *gmsweet15* mutant seeds also display defective endosperm function, characterized by the persistence of endosperm that is normally degraded in wild-type soybean seeds (Wang *et al*., 2019). This endosperm persistence in soybean suggests that SWEET-dependent sugar export is required not only for embryo growth but also for the proper developmental transition of the endosperm during seed maturation. In addition to having smaller embryos, the *zmsweet14a/14b/15a* mutant accumulates less oil in the embryo at maturity than wild-type (**Figure 3**). Although SWEETs are not directly involved in lipid or oil biosynthesis, the transported sugars likely provide carbon skeletons necessary for fatty acid synthesis (Rawsthorne, 2002), as illustrated in soybean (Miao *et al*., 2020; Wang *et al*., 2020, 2025; Zhang *et al*., 2020).

### ZmSWEET-dependent carbon partitioning and transcriptional regulation by DED1 at the embryo–endosperm interface

In addition to smaller embryos, the *zmsweet14a/14b/15a* mutant also shows reduced kernel weight, as has been reported for the single *zmsweet15a* mutant (Wang *et al*., 2024), as well as defects in carbon partitioning between the embryo and endosperm (**Figure 5**). One plausible explanation is that the mutations reduce sink strength at the embryo-endosperm interface, thereby reducing the overall carbon import into the two filial tissues. Alternatively, weak expression of *ZmSWEET15a* in the aleurone layer (Doll *et al*., 2020) could play a role in sugar import from maternal tissues (nucellus or integument) to the endosperm, thus directly limiting sugar entry into the endosperm. The overexpression of ZmSWEET15a was shown to increase the sucrose content in kernel, in agreement with a role of ZmSWEET15a in maintaining sucrose homeostasis in the seed (Liu *et al*., 2022). Interestingly, the R2R3-MYB transcription factor DOSAGE-EFFECT DEFECTIVE1 (DED1/ZmMYB73) has been identified as a direct regulator of these three EAS-expressed *ZmSWEET* genes targeted in our study (Dai *et al*., 2022). DED1 acts as a paternally derived regulator of kernel development and size in maize (Dai *et al*., 2022). Together with our findings, these observations suggest that modulation of EAS-expressed *ZmSWEET* transporters by DED1 represents one molecular mechanism by which DED1 influences kernel and embryo size, likely through the control of sucrose fluxes at the embryo-endosperm interface. However, the more severe kernel developmental defects reported in the *ded1* mutant compared with those observed in the *zmsweet14a/14b/15a* mutant indicate that DED1-mediated regulation of kernel development does not rely exclusively on ZmSWEET-dependent sugar transport, as supported by the fact that additional direct targets have been found for DED1 (Dai *et al*., 2022).

### Does sugar recycling in EAS cells link programmed cell death with embryo and endosperm growth?

Another intriguing aspect of EAS biology is the dynamic fate of the cell layers adjacent to the embryo scutellum, which are progressively eliminated as the embryo grows through a tightly regulated cell death process (Doll *et al*., 2020, 2025). This regulated cell elimination is associated with the remobilization of starch reserves (in the form of granules) accumulated before starchy endosperm cells acquire EAS identity(Doll *et al*., 2025). It is reasonable to hypothesize that the degradation products of starch in EAS cells must be efficiently exported and recycled before the cells die, a process that likely requires sucrose efflux mediated by ZmSWEET transporters. Consistent with this hypothesis, the altered carbon partitioning observed in both the embryo and endosperm of the *zmsweet14a/14b/15a* mutant (**Figure 5**) suggests that sugars released from EAS cells contribute to fueling two adjacent sink tissues; the rapidly growing embryo and the adjacent starchy endosperm, which remains a metabolic sink during maize kernel filling. Thus, ZmSWEET-mediated sugar recycling from EAS cells may represent a key coordination mechanism for carbon redistribution between competing sink tissues. Interestingly, maize kernels lacking an embryo (due to embryo defective *emb8522* mutation) show a strong ectopic activation of *ZmSWEET15a* gene expression in the scutellar aleurone layer (SAL), which is usually situated between the embryo and the aleurone (Doll *et al*., 2020). This suggests that the presence of embryo itself can modulate SWEET gene expression in the neighboring endosperm tissues, although the mechanism underlying this embryo-endosperm communication remains to be identified.

### ZmSWEET functions extend beyond seed filling to seedling establishment

In addition to seed development phenotypes, the *zmsweet14a/14b/15a* mutant also displays pronounced post-germinative defects, including reduced shoot and root fresh weight and impaired root development during early seedling growth (**Figure 4**). These phenotypes indicate that loss of *ZmSWEET14a/14b/15a* function impacts not only kernel filling but also seedling establishment. Two non-mutually exclusive hypotheses may account for these observations. First, the reduced embryo size combined with altered carbon accumulation during kernel development may result in embryos with diminished reserve availability and metabolic capacity. These “weaker” embryos could mobilize fewer resources during germination, ultimately leading to compromised seedling vigor compared with wild-type plants. Alternatively, ZmSWEET transporters may play a more direct role during germination and early seedling growth phases. In this scenario, the activation of SWEET-mediated sugar transport after imbibition would be required to sustain carbon allocation to rapidly growing tissues. This hypothesis is supported by studies reporting the transcriptional activation of at least 17 ZmSWEET genes in the primary root during germination, including the three ZmSWEET genes characterized in this study (Hoopes *et al*., 2019; López-Coria *et al*., 2019; Zhu *et al*., 2022) (**Supplemental figure S1B**). Together, these data support a dual role for ZmSWEET14a/14b/15a in both seed development and early seedling growth.

### Sucrose gradients and ZmSWEET function at the embryo–endosperm interface

Thanks to the development of FTIR imaging applied to maize kernel sections, this study reports, for the first time, the spatial imaging of sucrose distribution across the maize kernel. A clear sucrose gradient was detected within the endosperm, with higher sucrose abundance in the basal endosperm compared with the upper endosperm. Our imaging data are consistent with the recently metabolic compartmentation observed in the maize kernel (Chen *et al*., 2026). Sucrose was found to be highly enriched in the placento-chalazal region. This distribution is consistent with the current model of nutrient transfer into the maize kernel, in which assimilates are delivered primarily via phloem unloading at the pedicel. Nutrients released into the placento-chalazal region are then taken-up into endosperm via the BETL cell (Felker and Shannon, 1980; Patrick and Offler, 2001; Offler *et al*., 2003). Interestingly, sucrose imaging of maize kernels revealed additional sucrose accumulation in maternal tissues at sites other than the general buildup of sucrose in the pedicel. Indeed, both the tips of the kernel and the maternal tissues in contact with the developing embryo (**Figure 5**) accumulate sucrose. It will be interesting to investigate the proportion of sucrose suppied to the endosperm and the embryo through this route.

Taken together, our results support a model in which EAS-expressed ZmSWEET transporters play a central role in coordinating carbon transfer, recycling, and allocation at the embryo–endosperm interface. By regulating sucrose fluxes during seed development, these transporters influence embryo growth, kernel size and ultimately seedling vigor. Beyond their local transport function, the regulation of *ZmSWEET14a/14b/15a* expression requires integration of developmental programs (via DED1 signaling) as well as tissue-to-tissue communication, highlighting the EAS as a key regulatory hub for carbon partitioning in the maize kernel.

## Materials and methods

### Plant material and growth conditions

The maize (Zea mays) inbred line A188 and derived edited plants were grown under the French S2 safety standards for the culture of transgenic plants, in growth chambers or green house. The photoperiod consisted of 16 h light and 8 h darkness in a 24 h diurnal cycle. Temperature was set to 25°C/18°C (day/night). The relative humidity was controlled at 55% (day) and 65% (night). Seeds were germinated in 0.2 L of Favorit MP Godets substrate (Eriterre, Saint-André-de-Corcy) and transferred after 2 weeks to 8 L of Favorit Argile TM + 20% perlite substrate (Eriterre, Saint-André-de-Corcy) supplemented with liquid fertilizer. All plants were propagated by hand pollination.

### SWEET phylogeny

The 24 maize SWEET protein sequences were retrieved at from Gramene (http://www.gramene.org/) (Gupta *et al*., 2016), and the 17 Arabidopsis and 21 rice SWEET protein sequences were downloaded from Phytozome (https://phytozome.jgi.doe.gov/pz/portal.html). Conserved sites were selected using G-BLOCKS with all the three options available to minimize stringency. The phylogeny was built using PhyML, based on the amino acid alignment, as described previously (Guindon *et al*., 2009).

### Transient protoplast transformation for subcellular localization

#### Plasmid DNA Preparation

Plasmids are purified using the NucleoBond Xtra Midi/Maxi kit (Macherey-Nagel). Bacteria from glycerol stocks are plated on LB medium with 50 µg/mL spectinomycin. A single colony is used to seed 5 mL of LB with the same antibiotic and incubated at 37°C with agitation (250 rpm) for 8 h. This pre-culture is then expanded to 400 mL of LB containing spectinomycin and incubated overnight at 37°C, 250 rpm. Optical density at 600 nm is measured and adjusted to 4 before centrifugation and removal of the supernatant. Plasmid extraction follows the manufacturer’s protocol: cell lysis, neutralization, column loading, washing, elution, precipitation, washing, drying, and resuspension in sterile water. Plasmid quantity is assessed with spectrophotometry and NanoDrop, and quality is confirmed by restriction digestion and agarose gel electrophoresis, guided by SnapGene software predictions.

#### Protoplast Isolation and transformation

Protoplast Isolation protocol are close to previously published (Fierlej *et al*., 2022). A188 wild-type seedlings are grown in a greenhouse until emergence (∼0.5–1 cm) and transferred to darkness until the three-leaf stage (∼12 days after sowing). Leaf sections (∼1 mm) are cut from the central portion, avoiding the first leaf and leaf tips. Sections are incubated in 20 mL of enzyme solution (1.5% Cellulase R10, 0.75% Macerozyme R10, 0.1% Pectolyase Y23, 0.6 M mannitol, 10 mM MES pH 5.7, 10 mM CaCl_2_, 0.1% BSA) under vacuum (−500 mbar) for 30 min in the dark. Digestion continues 2 h at 25°C with gentle agitation (30 rpm). The enzyme-protoplast mixture is filtered through a 40 µm cell strainer, centrifuged at 100 g for 3 min (slow acceleration/deceleration), and washed twice with cooled 0.6 M D-mannitol. Protoplasts are finally resuspended in 2 mL of MMGZm buffer (0.6 M mannitol, 15 mM MgCl_2_, 4 mM MES, pH 5.7) and counted with a Malassez chamber. The concentration is adjusted to 1 × 10^6 protoplasts/mL and kept on ice in the dark. For protoplast transformation, 100 µL of protoplasts are mixed with 110 µL of freshly prepared PEG solution (40% PEG 4000, 0.2 M mannitol, 0.1 M CaCl_2_) and 20 µL of DNA (10 µg plasmid) in MMGZm. The mixture is incubated 10 min in the dark at room temperature, and the reaction is stopped with 800 µL of W5 buffer (154 mM NaCl, 125 mM CaCl_2_, 5 mM KCl, 2 mM MES, pH 5.7). Protoplasts are centrifuged at 100 g for 3 min, resuspended in 1 mL W5 buffer, and transferred to 24-well plates. Cultures are incubated in the dark at 25°C with gentle agitation (25 rpm) until further analysis.

#### Confocal microscopy

Protoplasts are imaged using an inverted confocal microscope (Zeiss LMS710) at 20 hours and 44 hours post-transformation. A 40X water-immersion objective (Plan-Apochromat 40×/1.0 water; Zeiss) is used. Fluorophores are excited with an argon laser at 458 nm for mTurquoise and 514 nm for mCitrine, while 561 nm excitation is used to detect autofluorescence. Images are acquired sequentially to separate channels according to their excitation and emission (Gilles *et al*., 2021). Detection settings are: 519-580 nm for mCitrine, 463-500 nm for mTurquoise and 600-795 nm for autofluorescence.

### Yeast heterologous complementation assay

To assess the sugar transport activity of the candidate genes, their full-length coding sequences were cloned into the yeast cDNA cloning vector pDR196, modified to be compatible with gateway technology pDR196-GW (Wipf *et al*., 2003; Koegel *et al*., 2013), and constructs were verified by sequencing. Recombinant pDR196-ZmSWEETs plasmids, together with the two type of empty vector pDR196 and pDR196-GW used as a negative control, were introduced into the yeast mutant strains SUSY7/ura3 (*Matα leu2-3 112 ura3-52 trp1 mal0 suc2::URA3 ura3 LEU2::PADH1-StSUSY1*), which is impaired sucrose uptake. SUSY7/ura3 cannot utilize extracellular sucrose unless a functional sucrose transporter is present at the plasma membrane (Riesmeier *et al*., 1992). Transformed yeast cells were cultured on synthetic deficient medium lacking uracil and supplemented with either 2% glucose or 2% filtered sucrose (Sigma Ultra, 99.5% GC ref: S7903-250) as the sole carbon source. Yeast cultures were adjusted to an OD_600_ of 0.1, serially diluted, and spotted onto solid media. Growth was monitored after incubation at 30 °C for 2 days for glucose containing medium and 3 days for sucrose containing medium. Functional complementation of sucrose uptake was evaluated based on the ability of the transformed strains to grow better as compared to empty vectors on sucrose containing medium.

### sgRNA design and CRISPR–Cas9 vector construction

Single guide RNAs were designed using the CRISPOR web tool (Concordet and Haeussler, 2018) based on the B73 maize reference genome and double check for compatibility with A188 genome. Two 20-nucleotide targets sequence were selected for each gene (Supplemental Table S1). Target sites were chosen based on high predicted on-target activity scores and the absence of predicted off-target sites with fewer than three mismatches in the maize genome. Because CRISPOR prioritizes guide specificity at the individual gene level, relatively low specificity scores were expected for targets shared among closely related gene family members. All genome editing-related cloning steps were performed exactly following the strategy described in (Doll *et al*., 2019). Briefly, the first sgRNA was cloned under the control of rice U3 promoter in binary vector L1609 containing a rice codon optimized Cas9 driven by constitutive maize ubiquitin promoter. The second sgRNA was cloned under the wheat U6 promoter in L1611 vector. The U6-driven gRNA target cassette from L1611 was subsequently excised with restriction enzymes and cloned into the L1609 derivative, downstream of the U3-driven target cassette, yielding a dual-sgRNA CRISPR construct targeting regions of the genes of interest (**Figure 2**). The two final binary vectors obtained (L1986 targeting ZmSWEET14a and ZmSWEET14b; L1987 targeting ZmSWEET15a) were verified by restriction analysis and Sanger sequencing prior to transformation. The validated constructs were introduced into Agrobacterium tumefaciens strain LBA4404(pSB1) by bacterial conjugation and used for maize transformation.

### Maize transformation and screen for edited plants

A. tumefaciens strain LBA4404 harboring the recombination product of pSB1 and the construct of interest was used to transform 12 DAP immature embryos of maize inbred line A188 according to a standard protocol (Ishida *et al*., 1996; Fierlej *et al*., 2022). Screening for genome editing events was performed on gDNA from T0 leaves by specific PCR amplification (Supplemental Table S1 for primer sequences) followed by Sanger sequencing. Chromatograms from wild-type and transgene-bearing plants were then compared to identify putative targeted mutagenesis events.

### MRI and TD NMR measurements

Non-invasive 3D measurements of maize (Zea mays) seeds were performed at 11.7 T Avance Neo 500 MHz Super Wide Bore using an NMR spectrometer (Bruker BioSpin) equipped with a 1H quadrature probe head with inner diameter of 66mm for multi-seed measurements (Plutenko *et al*., 2025). The seed holder was printed with a resolution of 50 μm on a Form 2 3D printer (Formlabs, USA) using high temperature non-magnetic resin (type flhtam01 and flhtam02). The MRI parameters for multiple seed screening were set as follows: field of view (FOV) 40×40×60mm, repetition time (TR) 750ms; echo time (TE) 3ms; average time 187 min and resolution 400 μm. For generating the water images, the pulses were applied resonant on the water frequency, and with an offset of −1400 Hz for the lipid images (Langer *et al*., 2023). Manual image segmentation was performed using Amira software (FEI Visualization Sciences Group, France). The lipid content was measured using TD-NMR (MQ60 instrument, Bruker, Germany) as detailed earlier (Rolletschek *et al*., 2015).

### Sucrose mapping using high resolution FTIR

Quantitative mapping of sucrose distribution in the maize caryopsis was done using Fourier Transform Infrared microspectroscopy (FTIR). Imaging and data processing were done as essentially described in (Guendel *et al*., 2018). In short, intact caryopses were harvested at 13 DAP, instantly frozen in liquid nitrogen, embedded in Tissue-Tek cryomolds at −20°C, and cross sectioned (6 µm) with a cryotome (CryoStar NX7, ThermoFisher Scientific). Tissue sections were lyophilized and stored in darkness at room temperature until analysis. Imaging was performed using a Hyperion 3000 FTIR microscope (Bruker Optics) coupled to a Tensor 27 FTIR spectrometer (Bruker Optics) with an internal mid-infrared source. FTIR images were recorded in the spectral range of 3900 cm^−1^ to 800 cm^−1^ at a spatial resolution of 11 µm and a spectral resolution of 6 cm^−1^. Data processing and sucrose fingerprinting was done using software MatLab (The MathWorks) exactly as described.

### Germination in hydroponic assay

Maize seeds were surface-sterilized in 1.4% (v/v) bleach, 1 ‰ (v/v) Tween-20, for 15 min under gentle agitation. The seeds were then treated with H2O2 (35%; v/v) for 2 min, rinsed with 70% (v/v) ethanol, and washed 6 times with sterilized water. The seeds were overlaid with wet clay beads in a plastic box, which was itself covered by a transparent plastic film. Seeds were germinated and further grown in a growth chamber at 65% relative humidity, with 15 h/9 h light/dark cycles (150 μE m-2 s-1) at 22°C (light)/20°C (dark). At 5 DAS, seedlings were transferred to hydroponic containers (61.6 X 35.8 X 13.6 cm; 20 seedlings /container) filled with 24 L of a medium containing 1.25mM KNO3, 0.75mM MgSO4, 1.5mM Ca(NO3)2, 0.5mM KH2PO4, 0.1mM MgCl2, 0.05mM Fe-EDTA, 0.05mM H3BO3,0.012mM MnSO4, 0.7mM CuSO4, 0.001mM ZnSO4, 24.10-5Mm MoO4Na2,1.10-5 mM CoCl2, 0.1mM Na2SiO3, and 1mM MES. Plants were grown in these solution for 6 days. At 11 DAS, roots were excised and imaged for their root system architecture. Root and shoots were weighed.

### Isotopic mass spectrometry

13DAP Kernels of *zmsweet14a/14b/15a* and wild type are compared for their accumulation 13C-labeklled sucrose on all carbons (CLM-7757-PK, CAS #: 41055-68-9, Cambridge Isotope Laboratories, Inc. (USA)). The kernels are separated from the ear using a scalpel blade, taking care not to damage them and keeping a pedicel length of approximately 2 mm. The glumes are removed and the kernels are rinsed 3-4 times for 30 seconds each in 1X PBS, then the end of the pedicel is trimmed by approximately 0.5 mm with a fine razor blade. Kernels are then incubated in a 96-well plate containing 2% (w/v) labeled sucrose, diluted in 1X PBS. The ^13^C-sucrose solutions are prepared as follows: a final 2% sucrose solution containing 10% ^13^C sucrose diluted in 1X PBS. The solution is then dispensed at 160 µl per well. In order to prevent evaporation, the plates are placed at room temperature, and 48 kernels per replicate are collected and dissected at the indicated times. During sampling, the kernels are rinsed in two successive baths of sterile 1X PBS (∼10 ml), followed by a final rinse with 1X PBS to remove non-accumulated ^13^C. To dissect the different parts of the kernel, the pericarp is incised on either side of the embryo and then removed (Doll *et al*., 2020). The embryo is removed, rinsed in sterile 1X PBS, and briefly blotted on absorbent paper for 5 seconds, then placed in an aluminum cupule. The endosperm is cut transversely just above the space left by the embryo, resulting in an upper and a lower part. These parts are placed in separate tin cupules. Finally, the samples are dehydrated by placing them in a ventilated oven for 4-5 days with the following settings: Temperature: 60°C; Fan: 70%; Valve: 50%. Samples are analyzed by Isotopic mass spectrometry (Vario-PYROcube elemental analyzer coupled with an IsoPrime Precision mass spectrometer, Elementar, UK) by the AQuI plateform (Montpellier, France). The dried and ground samples are placed in a tin cupule and injected into an elemental analyzer (Vario-PYROcube, Elementar, UK). After combustion at 920°C in the presence of oxygen and CuO, the gas molecules from the sample (mainly H_2_O, CO_2_, N_2_, and N_2_O) are transported via a carrier gas flow (ultra-pure helium) to a reduction furnace where the nitrogen oxides are reduced to N_2_ in the presence of copper at 600°C (Dumas reaction). The H_2_O produced is trapped by SICAPENT columns (Merck). The N_2_ and CO_2_ are then separated. The CO_2_ is trapped at room temperature in a TPD (Temperature Programmable Desorption) column and subsequently released at 100°C. A thermal conductivity detector (TCD) quantifies total N and C. N_2_ and then CO_2_ are then analyzed by mass spectrometry (IsoPrime Precision, Elementar, UK) to determine the isotopic abundances ^13^C/ ^12^C.

The δ ^13^C is calculated using the following formula and has no unit:

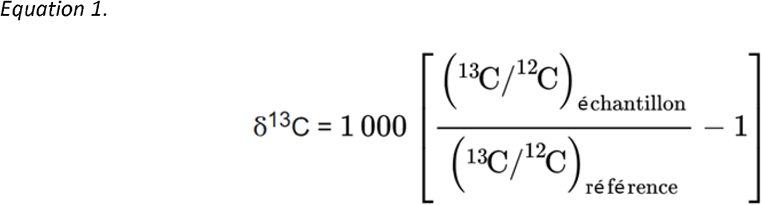

The reference is Pee Dee Belemnite, the shell of a fossil belemnite from the Cretaceous period. This reference has an abnormally high ^13^C/^12^C ratio (0.0112372), meaning that most natural samples have a negative δ^13^C (Slater *et al*., 2001). Statistical and graphical analyses were produced using R studio software. The primary objective was to establish a relationship between the measured parameter, δ^13^C, and the factors tested, such as genotype, compartment, and soaking time.

## Acknowledgements

The authors are thankful to Camille Knaupp, Justin Berger, Patrice Bolland, Isabelle Desbouchages, and Hervé Leyral for technical assistance in maize culture and media preparation; Cindy Vial, Laureen Grangier, Nelly Camilleri, and Julie Prata for administrative assistance. We also thank I. Plutenko, S. Wagner and S. Ortleb for experimental support in seed phenotyping and MRI. We acknowledge Joan Doidy for providing the SUS7 yeast strain and the associated yeast protocols together with Gael Yvert. HR and LB acknowledge funding by the Federal Ministry of Education and Research (MAGDI-Project; BMBF grant 031B1451A), the Deutsche Forschungsgemeinschaft (grant no. RO 2411/7-2), the European Regional Development Fund (ERDF) and the Investment Bank of Saxony-Anhalt (FKZ: ZS/2019/09/101444). YF was supported by a CIFRE fellowship of the ANRT (grant N°2018/0480). LG was supported by a Ph.D. fellowship from the Ministère de l’Enseignement Supérieur et de la Recherche. TW and BL acknowledge support from the Plant Science and Breeding Division of the Institut National de la Recherche en Agriculture et Alimentation et Environnement (BAP, INRAE). We acknowledge the contribution of SFR Biosciences (Universite Claude Bernard Lyon 1, CNRS UAR3444, Inserm US8, ENS de Lyon) for support in imagining and the help of the staff of LyMIC-PLATIM. We honor the memory of Peter Rogowsky, an irreplaceable colleague, without whom this “*SWEET manuscript*” would not exist.

