## Supplemental figure S1 for "ZmSWEET Sucrose transporters expressed in the endosperm adjacent to the maize embryo are necessary for carbon partitioning and embryo growth"

**A**

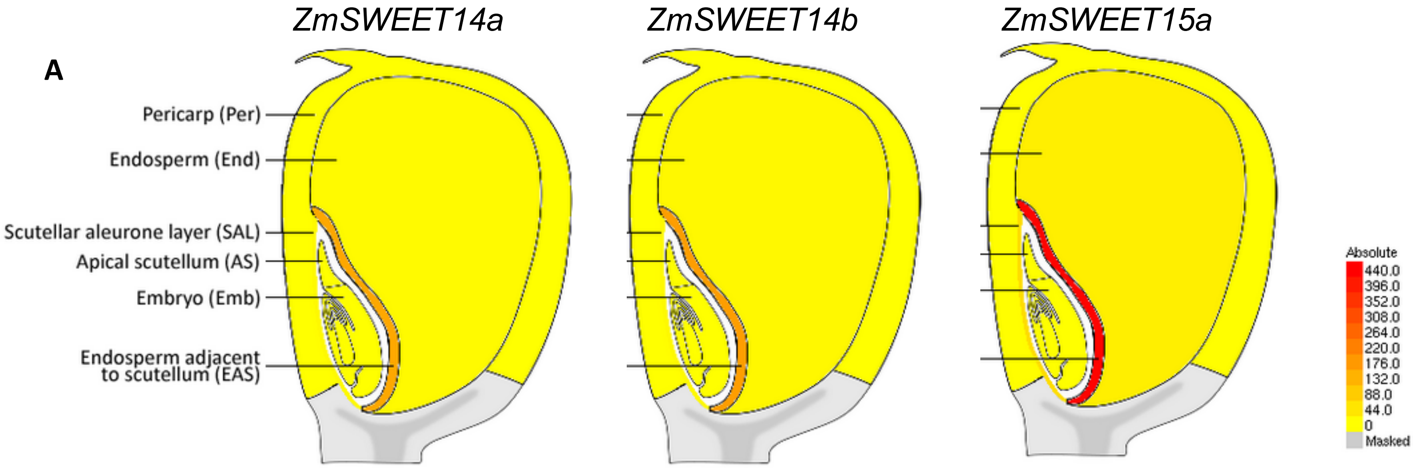
ZmSWEET14a
ZmSWEET14b
ZmSWEET15a
Pericarp (Per)
Endosperm (End)
Scutellar aleurone layer (SAL)
Apical scutellum (AS)
Embryo (Emb)
Endosperm adjacent to scutellum (EAS)
Absolute
440.0
396.0
352.0
308.0
264.0
220.0
176.0
132.0
88.0
44.0
0
Masked

**B**

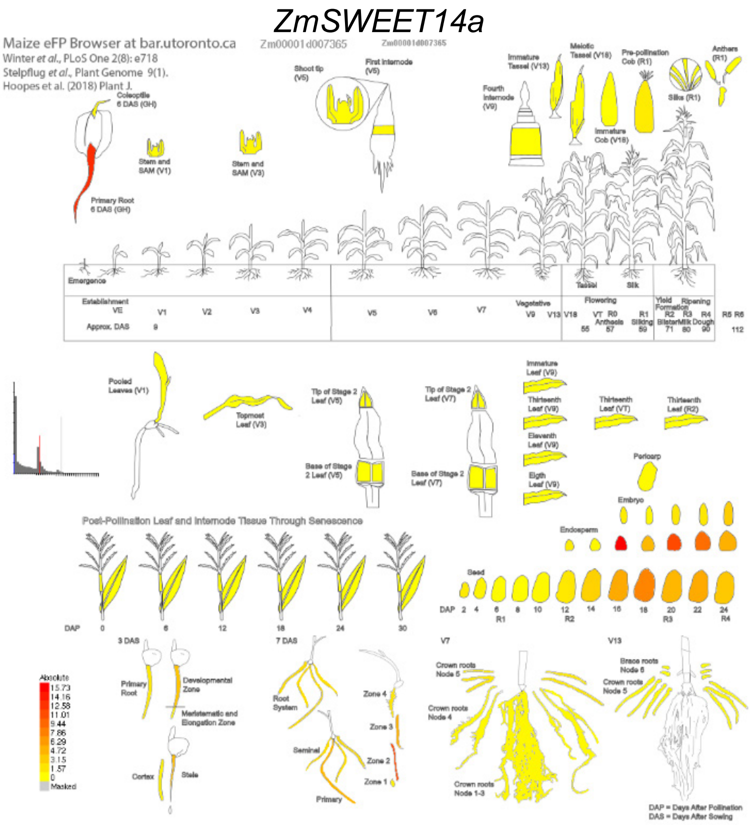
ZmSWEET14a
Maize eFP Browser at bar.utoronto.ca
Zm00001d007365
Winter et al., PLoS One 2(8): e718
Stelpflug et al., Plant Genome 9(1).
Hoopes et al. (2018) Plant J.
Post-Pollination Leaf and Internode Tissue Through Senescence
DAP = Days After Pollination
DAS = Days After Sowing

*ZmSWEET14b*

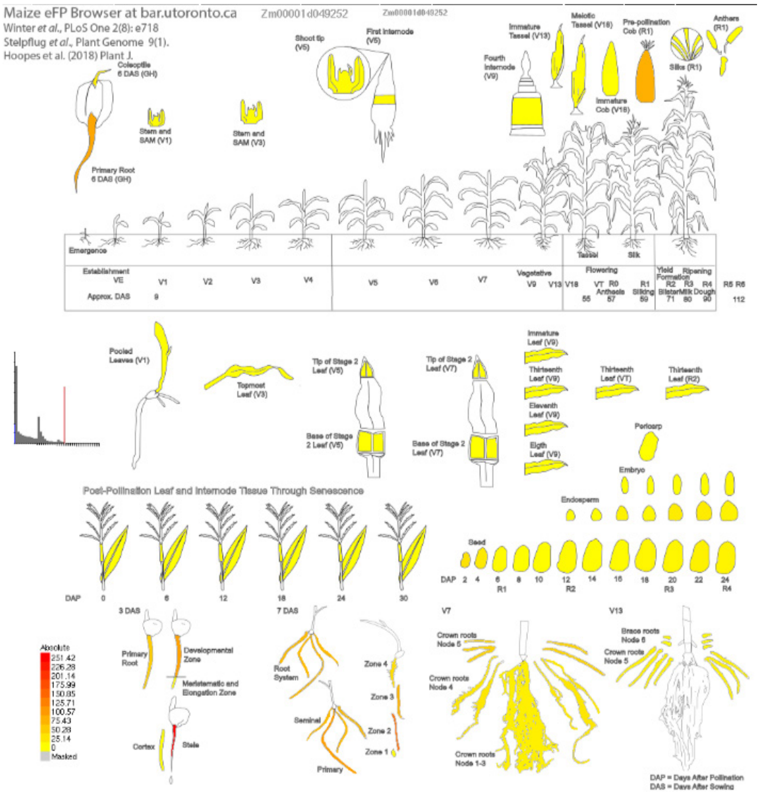


*ZmSWEET15a*

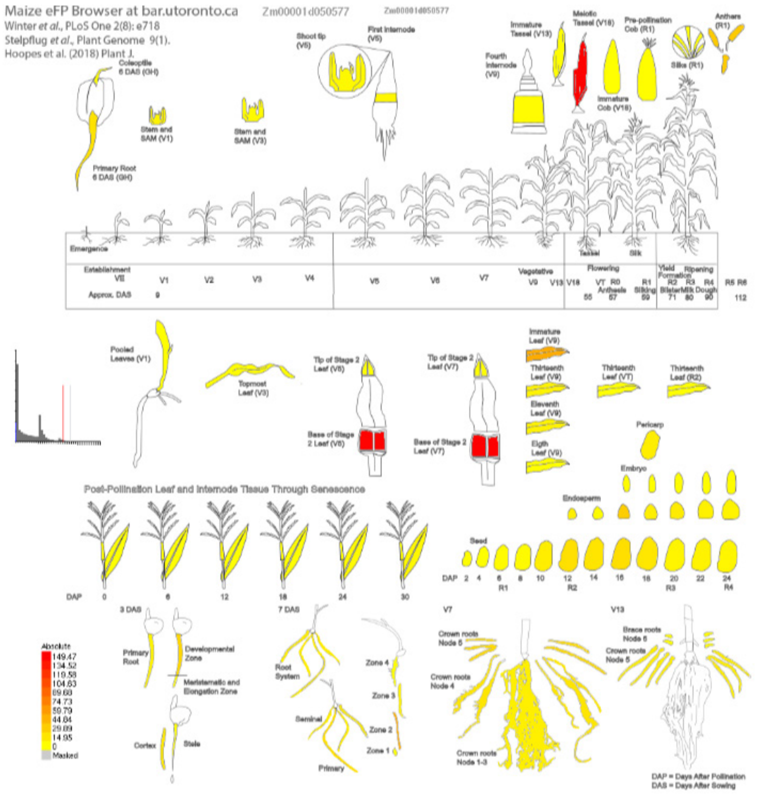


**Supplemental figure S1. Expression profiles of *ZmSWEET14a/14b/15a* using Maize eFP Browser**

**A)** Expression profile in 13 DAP-old kernels using doll et al., (2020) data set. Signal threshold were set up similarly for all three genes to be comparable.
**B)** Expression profile in different maize tissues using Hoopes et al., 2019 data set.
