## Supplemental figure S2 for "ZmSWEET Sucrose transporters expressed in the endosperm adjacent to the maize embryo are necessary for carbon partitioning and embryo growth"

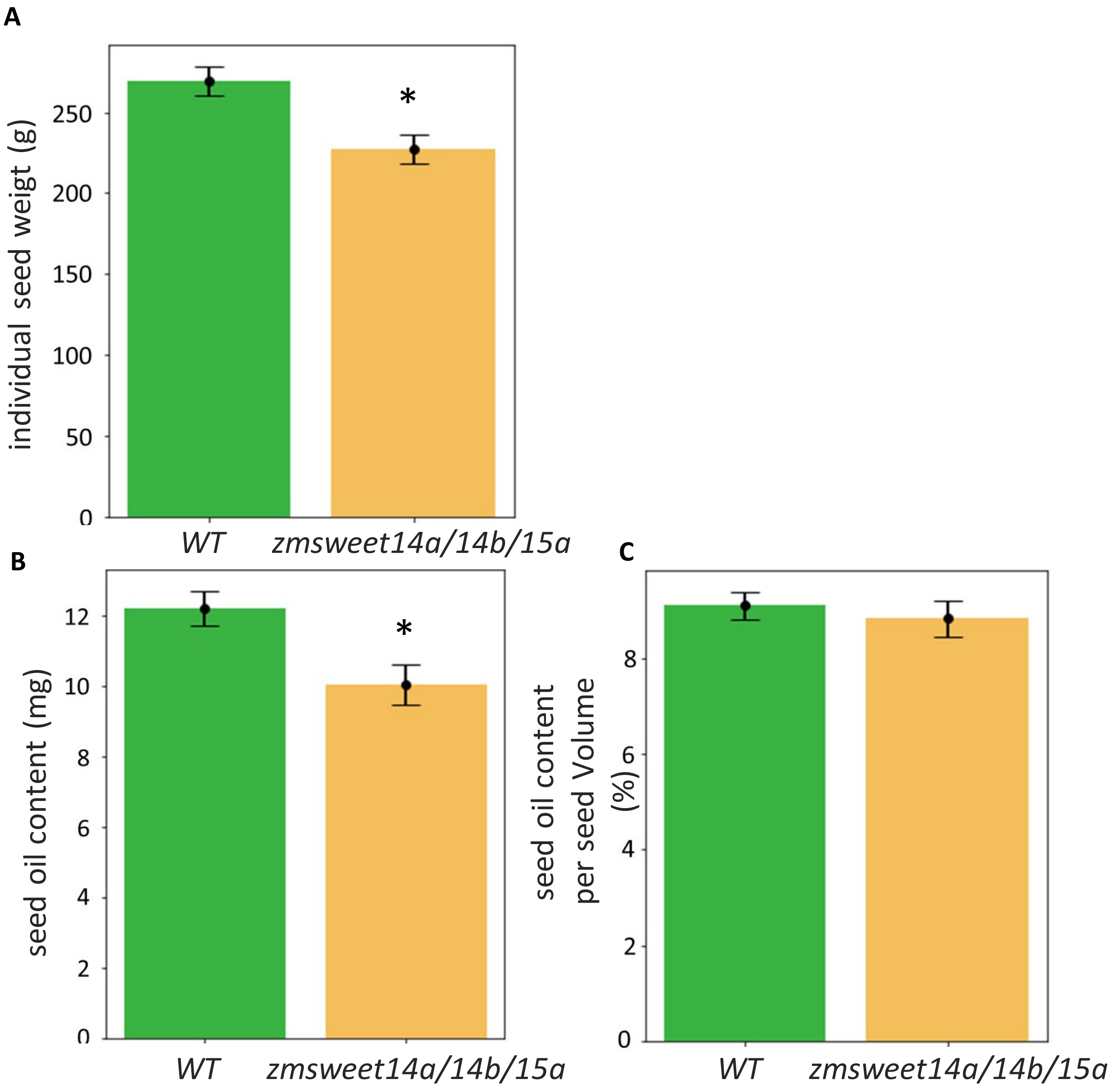


**<u>Supplemental figure S2</u>. Reduced seed weight and oil content in the *zmsweet14a/14b/15a* mutant line.**

**(A)** Mean individual seed weight (n = 40 seeds per genotype).
**(B)** Mean seed oil content expressed as milligrams per seed (n = 40 seeds per genotype)
(**C**) Percentage of oil content measured using time-domain nuclear magnetic resonance (TD-NMR). It represents the relative concentration of oil within the seed volume (n = 40 seeds per genotype). * Asterisks indicate statistically significant differences ($p<0.05$, n=40, T-test).
